# From Metapopulation to Metacommunity and the Complications of Community Structure

**DOI:** 10.1101/504670

**Authors:** Zachary Hajian-Forooshani, Lauren Schmitt, Nicholas Medina, John Vandermeer

## Abstract

The metacommunity, as it evolved from Levin’s metapopulation, provides a framework to consider the spatial organization of species interactions. Arguably the most fundamental feature of metapopulations and metacommunities are that demes are connected via migration. An important result from Levin’s metapopulation work—that increasing migration lowers regional extinction probability—is often incorporated into conceptions of metacommunities. We first use a toy model to show how this result from Levin’s metapopulation does not necessarily hold when considering community interactions in a metacommunity context. We also report results from a metacommunity field experiment conducted with a tropical terrestrial leaf litter community and show that migration induces the extinction of predators in the metacommunity. Our result corroborates the findings of a prior metacommunity experiment in a temperate terrestrial leaf litter community. The concordance between these experiments even with vastly different communities highlights the importance of considering trophic and non-trophic community structure to understand metacommunity dynamics.

## 1 Introduction

The theory of metapopulations has become a standard way of thinking about simple population dynamics, and its success has stimulated a seemingly obvious extension, the metacommunity (Levins 1969; Wilson 1992). As originally envisioned, a metacommunity is a collection of interacting populations in which extinction and migration occurs on a regular basis. This in turn results in patchiness that creates subcommunities which may be distinct in species composition from one another. It might be argued that MacArthur and Wilson’s theory of island biogeography was the first metacommunity theory, and perhaps the most elegant, in which the patchiness is provided by the existence of islands (MacArthur and Wilson 1964). However, an important conclusion of metapopulation theory has been tacitly incorporated as a clear and obvious corollary—that increasing migration lowers overall extinction probability. While it may be a reasonable proposition at first glance, further reflection on the assumption suggests that the lowered extinction expectation is not universal in the metacommunity context (Vandermeer et al. 1980; Caswell and Cohen 1991).

Huffaker’s classic experiment might be thought of as a canonical case study that supports a pervasive link between migration and lower extinction. A predator/prey combination, when isolated in a physically small space, goes extinct (predator eats prey to extinction then itself goes locally extinct), yet when extended in space through prey and predator migrations, apparently stable oscillations result (Huffaker 1958). The same pattern was observed in Gause’s experimental system of protozoans twenty years prior (Gause et al. 1936). Alternatively, however, it is not difficult to imagine the reverse situation: a case in which migration might increase extinction probability. For example, in a two-predator one-prey situation in which spatial structure allows for a segregation of the two predators in space; increasing the predator migration rate could increase intraguild antagonism, leading to one of the predators dominating and a reduction of species diversity from three to two. Thus, elevated migration levels could result in either increased or decreased species diversity, depending on the strength of antagonistic (or even facultative) interactions across ecological guilds. The conclusion from this must be that the simple migration/extinction equilibrium of island biogeography and metapopulation theory may be too simple for more complex community structures.

A simple toy model illustrates this point. Consider a contextual division of a community into p, a species affected interspecifically by q, a species which is not affected by p. In the spirit of the Levins metapopulation concept, p and q represent fractions of habitats occupied by each species. For example, p could be a prey species and q a predator species, or q could be a species that exerts a behavioral modification on p such that its local extinction rate is increased through a second order effect. Other well-known relations could be cited (Werner and Peacor 2003; Terry et al. 2017). In a metacommunity context we simply note that q increases the extinction rate of p. The simplest model possible would be:

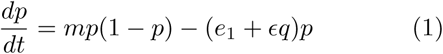

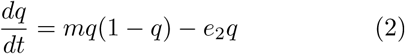

where m is migration rate, *e*_*i*_ is the base extinction rate for *p* and *q*, *ϵ* is the added extinction factor from species *q*. Equilibrium values of both *p* and *q* are displayed in Figure 1, which shows that a reduction in the equilibrium value of *p* with increased migration is possible for a range of values of *m*,. Whether equilibrium values increase or decrease as a function of increasing migration depends on the added extinction factor *ϵ* —when *ϵ* is high, extinction of an affected population (represented here by *p*) is possible with increased migration.

**Figure 1:**
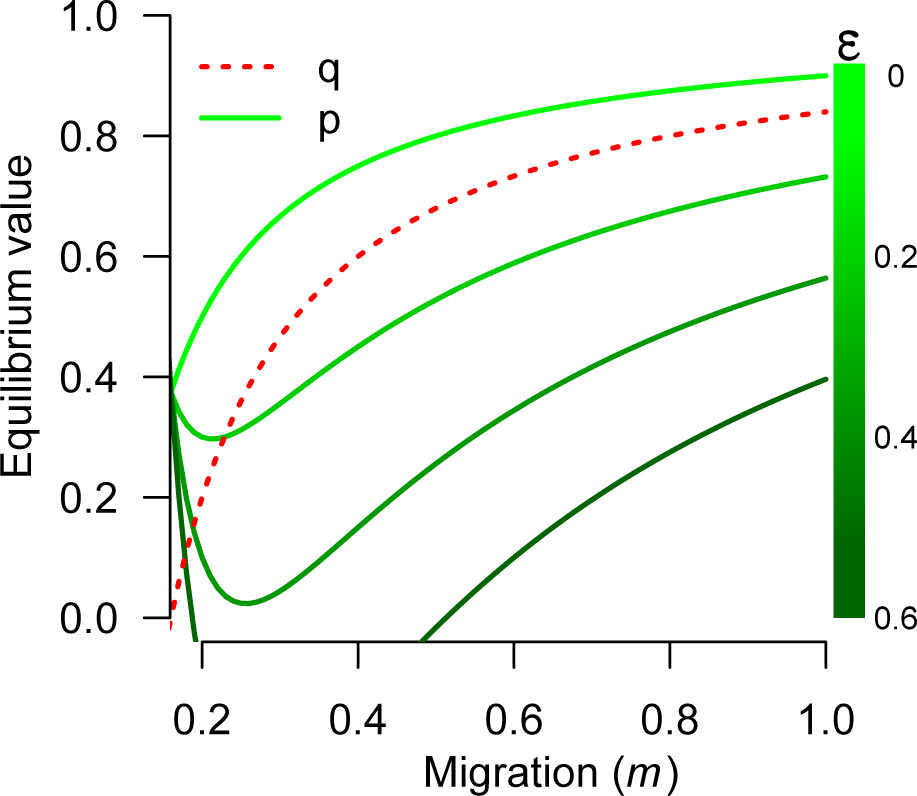
Equilibrium values of *p* (non-monotonic green curves, with *ϵ* scale on right) and *q* (monotonic increasing red curve) from the basic toy model, illustrating how the indirect effect of *q* on *p* alters the equilibrium value of *p* (base parameters are *e*_1_ = 0.10; *e*_2_ = 0.16). Analytical details in supplementary material.

Others have highlighted that the role of migration in rescuing unstable community elements often seems to have been taken as a generalization, but it is evident (see Equation 1 and Equation 2) that the result is not always theoretically inevitable (Simberloff and Cox 1987). The ability to recreate a wide range of metacommunity dynamics theoretically suggests the need for an empirical approach. One of the key shortfalls of much of the experimental metacommunity work lies in its simplification of community interactions (Polis et al. 1989). Often only a subset of real communities frequently structured by trophic interactions are included (Warren 1996; Shurin 2001; Kneitel and Miller 2003; Cadotte 2006; Fox et al. 2017). Although there are some notable experiments which attempt to include some of the trophic and non-trophic realism of communities (Neill 1974; Vandermeer et al. 1980), surprisingly few experimental studies have focused on such empirically realistic metacommunities.

One of the early attempts to study the role of migration in empirical metacommunities used leaflitter macro-arthropod communities and found that the predator guild (defined taxonomically) actually decreased in richness when random migration was induced while non-predator richness was unaffected by migration (Vandermeer et al. 1980). These results suggest that generalizations about metacommunity structure may be trophic-specific at times. This idea is already implicit in some well-known debates, for example, top-down versus bottom-up control (Hairston et al. 1960) or trophic cascades operating differently in aquatic versus terrestrial systems (Strong 1992). A wide range of ecological processes likely interact with migration in real metacommunities, such as higher-order effects which recent work suggests may be more determinant of community structure than the more direct, lower-order effects (Werner and Peacor 2003; Bairey et al. 2016; Grilli et al. 2017; Terry et al. 2017).

The empirical result that experimental migrations make a difference in metacommunity structure for predators, but not for non-predators (Vandermeer et al. 1980), was found in a species-poor temperate deciduous forest (Michigan, USA). As a test of these results’ robustness across highly divergent ecosystems, it is of interest to query whether similar results might be obtained in a more speciose tropical ecosystem. It could be that the vaunted stability-preserving aspect of high biodiversity could overwhelm any special effect of strongly antagonistic predator species, which was suggested to have happened in the earlier temperate zone study. Here we revisit the experiment conducted by Vandermeer et al. (1980), but this time in a montane tropical agroecosystem. Accordingly, we sought to investigate how migration affects leaf-litter community richness and composition, focusing especially on the effects at different trophic levels. Based on underlying assumptions of the importance of migration in biodiversity, we hypothesized that curtailing local migration would reduce local species diversity, and that this effect would be observed at all trophic levels.

## 2 Methods

### 2.1 Study region and design

This study was conducted at Finca Irlanda, an organic shaded coffee agroecosystem in the Soconusco region of Chiapas, Mexico. The study site was on land recently transitioned from rustic coffee production to a forested reserve. The experimental set-up was positioned adjacent to a patch of invasive golden bamboo, the litter of which created a uniform mat. Leaf litter was collected from a well-forested area of the reserve, homogenized, and separated into 10 mesocosms. Mesocosms were 0.5 m2 and separated from one another by 1 meter. Five mesocosms were positioned on either side of a walking trail. No physical barriers prevented migration between mesocosms; we assumed organisms would be unlikely to leave a mesocosm of leaf litter to migrate across a relatively unhospitable mat of dried bamboo litter.

Half of the mesocosms were assigned as controls and half as treatments. Every four days for 16 days one-quarter of the litter was “migrated” between treatment mesocosms. The transfer pattern was randomized and a different quarter of the mesocosm was migrated during each transfer event. No migration took place between control mesocosms, though one-quarter of the litter was removed, agitated and replaced in the same mesocosm every 4 days to control for the disturbance of the litter transfers among treatment mesocosms. All mesocosms were harvested on day 20.

The litter was sieved using 3-mm meshes to remove large detritus. The sample was searched by four people for 20 minutes, and all encountered organisms were individually removed and placed in alcohol. This technique did not likely capture all organisms found within the mesocosms, but we expect that any bias toward certain groups of organisms was standardized across all samples, which we assured by blinding the sample labels throughout the sorting process. Individuals were sorted into orders or families and identified to morphospecies. Morphospecies were then classified as either predators or non-predators, where predators included spiders, Staphylinidae beetle larvae, pseudo-scorpions and centipedes.

### 2.2 Statistical methods

To look at the number of species in our control and migration treatments, individual-based rarefaction curves were calculated for the whole data set then separated into the two trophic groups of predators and non-predators. Rarefactions followed the now standard methodology of resampling the list of species observations with replacement at increasing numbers of individuals (Gotelli and Colwell 2001). One thousand resamples were conducted for each level of individuals sampled, and the mean number of species for a given density of individuals was calculated.

While there are standard methods to extrapolate the number of species (Colwell and Coddington 1994; Chao et al. 2014), and to compare the overall shape of rarefaction curves (Cayuela et al. 2015), we were interested in understanding how our curves differed along the axis of individuals sampled. To do this, we implemented a bootstrapping procedure to compare the difference between the number of species in the control and migration treatment compared to the number of species of a pooled data set for a given level of individuals sampled. This was done by resampling both control and migration datasets and calculating the difference in the number of species observed (Δ_*OBS*_ = *S*_*c*_ – *S*_*e*_) where *S*_*i*_ is the average number of species obtained in 1000 random draws from the actual species pool in the *i*^*th*^ treatment (either *c* or *e*). We then pooled both control and migration data together and randomly partitioned the data into two groups corresponding to the number of individuals obtained in the experimental data, resulting in artificial species lists corresponding to the number of individuals in the experimental control and migration data sets. We resampled both control and migration artificial datasets and calculated the difference in the number of species observed, here referred to as 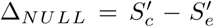, where 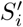 is the average number of species obtained in 1000 random draws from the actual species pool in the *i*^*th*^ treatment (either artificial *c* or artificial *e*). If Δ_*NULL*_ *>* If Δ_*OBS*_ we set *p*_*i*_ = 1, otherwise *p*_*i*_ = 0. Repeating this procedure 200 times, we calculated the significance probability as 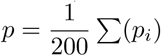.

## 3. Results

While there was no statistically significant difference between the control and migration treatments for neither the whole community (Figure 2A) nor the non-predator community (Figure 2C), we did observe a significant difference in species richness between the control and migration treatment within the predator community (Figure 2B). This difference between control and migration treatments for the predators starts at just nine individuals sampled and remains significant for the rest of the overlap between the two curves shown.

**Figure 2:**
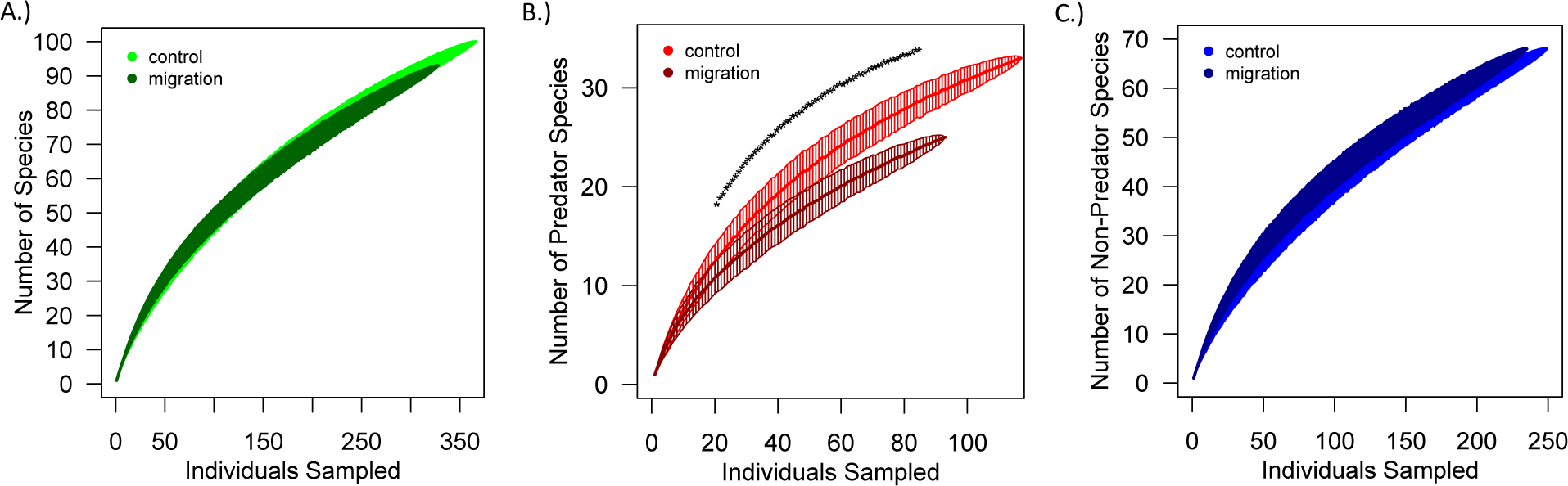
(A-C)Individual-based rarefaction curves for A.) whole community (green), B.) predators (red), and C.) non-predators (blue). Treatments are shown in lighter colors (control) and darker colors (migration). One standard deviation (based on the 200 random draws) is plotted in the shaded areas around the curves. above the curves represent a statistically significant (*p* < 0.05) difference in the number of species for a given number of individuals sampled between the control and migration treatment.

## 4. Discussion

While theoretical treatments of metacommunities acknowledge the potential complexity of community structure and its effect on migration (Caswell and Cohen 1991; Mouquet and Loreau 2002; Economo and Keitt 2008), it remains that simplified metacommunity theory generates the prediction that migration will tend to cause species diversity to increase, a result in concordance with the original MacArthur/Wilson, Levins/Heatwole framework (Heatwole and Levins 1972; Heatwole and Levins 1973; Levins et al. 1973). Contrarily, our results repeat those of the earlier study (Vandermeer et al. 1980) in which the predator guild species diversity decreases as migration increases, even after scaling up the low biodiversity background used in most experimental metacommunity studies (including our previous study in a temperate zone leaf litter community) to a tropical leaf litter community with over 100 different species. It is notable that even the coarsest distinction of trophic complexity (predators and non-predators) provides insights that do not emerge when analyzing the community as a whole. Importantly, there is also no evidence, to our knowledge, that suggests that leaf litter communities in the temperate or tropical zones are organized in such a way that predisposes them to the results of both of these studies. These consistent results with distinct communities in distinct regions suggest that there may be some generality in the way that migration impacts trophic structures.

We concur with our earlier study and hypothesize that a highly antagonistic and relatively rare (possibly initially isolated to only a single patch) predator may be shaping the community when dispersed among patches (Vandermeer et al. 1980). Given the relatively short timescale of our experiment (3 weeks), it also seems possible that the changes observed may result from non-trophic interactions (Werner and Peacor 2003; Bairey et al. 2016; Grilli et al. 2017; Terry et al. 2017). Anti-predator behaviors have been shown to shape metacommunity dynamics when migration is induced in simple experimental systems (Kneitel and Miller 2003; Hauzy et al. 2007; Howeth and Leibold 2010), but most studies use simplified versions of ecological communities that likely do not incorporate the trophic and non-trophic features captured by the experimental system reported here.

The work reported herein sits comfortably with the current enthusiasm for metacommunities, a framework originally suggested by Wilson (1992). It is substantially similar to the framework of MacArthur and Wilson’s original offering, in which 1) ecological dynamics occur locally, with species interactions of various forms determining which species will survive and which of those will perish (locally), while 2) the more regional process of migration continually feeds these local communities, countering local extinctions with regional migrations to provide the expected equilibrium (MacArthur and Wilson 1964). Eschewing some recent complexities (Leibold et al. 2004), we consider a metacommunity as structured in the original sense of Wilson (1992), wherein ecological dynamics occur at a local level, but local patches affect one another through dispersal. Our experiment seeks to explore the consequences of migration, but more specifically explores the interaction between community structure and the dynamics of migration.

## 6 Appendix

The basic model,

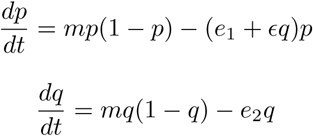

has evident equilibirum points at,

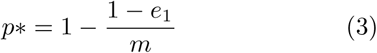

and

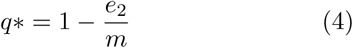

Substituting 4 into 3,

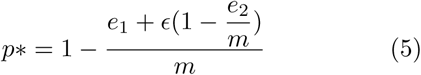

Whence it is clear that

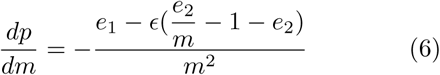

Which will be negative whenever,

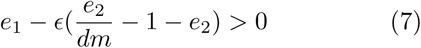

or

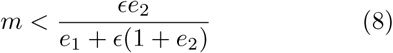

Note that from equation 1, the equilibrium value of p will be negative (i.e., extinction) if

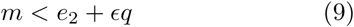

which, when substituting q* for q we have,

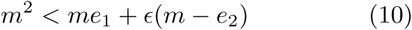

whence,

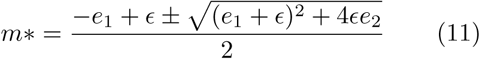

brackets the range of *m* for which the *p* species is extinct.

